# Modulation of Yorkie activity by alternative splicing is required for developmental stability

**DOI:** 10.1101/2019.12.19.882779

**Authors:** Diwas Srivastava, Marion de Toledo, Laurent Manchon, Jamal Tazi, François Juge

## Abstract

The mechanisms that contribute to developmental stability are barely known. Here we show that alternative splicing of *yorkie* (*yki*) is required for developmental stability in *Drosophila*. Yki encodes the effector of the Hippo pathway that has a central role in controlling organ growth and regeneration. We identify the splicing factor B52 as necessary for inclusion of *yki* alternative exon 3 that encodes one of the two WW domains of Yki protein. B52 depletion favors expression of Yki1 isoform carrying a single WW domain, and reduces growth in part through modulation of *yki* alternative splicing. Compared to the canonical Yki2 isoform containing two WW domains, Yki1 isoform has reduced transcriptional and growth-promoting activities, decreased binding to PPxY-containing partners, and lacks the ability to bridge two proteins containing PPxY motifs. Yet, Yki1 and Yki2 interact similarly with transcription factors and can thus compete *in vivo*. Strikingly, flies deprived from Yki1 isoform exhibit increased fluctuating wing asymmetry, a signal of increased developmental noise. Our results identify *yki* alternative splicing as a new level of control of the Hippo pathway and provide the first experimental evidence that alternative splicing participates in developmental robustness.

## Introduction

The Hippo signaling pathway is a conserved pathway that regulates cell fate and cell proliferation to control organ growth and regeneration ^1^. The core of the pathway comprises a kinase cascade including Hippo (Hpo, MST1/2 in mammals) and Warts (Wts, LATS1/2 in mammals), which phosphorylate the transcriptional coactivator Yorkie (Yki, YAP in mammals) to sequester it in the cytoplasm through binding to 14.3.3 proteins. Upon inactivation of the pathway, unphosphorylated Yki/YAP translocates into the nucleus and activates a transcriptional program promoting cell proliferation and inhibiting apoptosis. Yki/YAP does not bind DNA directly but interacts with specific transcription factors, especially members of the TEAD family Scalloped (Sd) in flies and TEAD1/2/3/4 in mammals. Nuclear Yki/YAP then recruits the histone methyl-transferase NcoA6 to trigger transcription activation ^2,3^. In addition, Yki was proposed to displace the transcriptional repressor Tgi (Tondu-domain-containing Growth Inhibitor, SDBP in mammals) from Sd/TEAD and relieve a transcriptional repression ^4,5^. Yki/YAP interacts with its partners through different domains: the N-terminal domain binds Sd/TEAD whereas the two WW domains mediate interaction with PPxY motifs-containing proteins such as the repressor Tgi and the transactivator NcoA6 for example. Interestingly, both Yki and YAP exist as two isoforms containing either one or two WW domains, owing to the alternative splicing of an exon encoding the second WW domain ^6,7^. In mammals, YAP isoform with only one WW domain cannot bind angiomotin or p73 indicating that alternative splicing could play a role in the modulation of YAP activity ^8,9^. Up to now, the molecular bases and as well as the functional importance of this alternative splicing event *in vivo* are unknown.

Here we identified *yki* as an alternative splicing target of the splicing factor B52 in *Drosophila*, and functionally address the importance of *yki* alternative splicing *in vivo*. The B52 protein belongs to the family of Serine and arginine-rich (SR) proteins that are conserved RNA binding proteins involved in several steps of mRNA metabolisms and which play a major role in both constitutive and alternative splicing ^10^. We previously showed that, in *Drosophila*, the level of B52 protein influences cell growth ^11^, but the underlying molecular mechanisms were not characterized. Here we show that B52 is necessary for inclusion of *yki* alternative exon and that B52’s depletion reduces growth in part through modulation of *yki* alternative splicing. We further show that alternative inclusion of Yki second WW domain is an additional level of modulation of Yki activity that is unexpectedly required for correct growth equilibrium between right and left sides of *Drosophila*. These results identify the first alternative splicing event involved in developmental robustness.

## Results

### *yki* alternative splicing is a target of B52

We previously observed that B52 depletion reduces cell growth in *Drosophila* larvae salivary glands as well as in the thorax at later stages ^11^. We reasoned that B52’s effect on growth might be due to alternative splicing modulation of one or several genes controlling growth. To identify splicing events that are robustly affected by B52 depletion, we crossed two previously published RNA-seq datasets corresponding to RNAi-mediated depletion of B52 in S2 cells compared to control cells ^12,13^. We used MAJIQ (Modeling Alternative Junction Inclusion Quantification) ^14^ to identify Local Splicing Variations (LSV) that vary more than 20% between control and B52 RNAi conditions, in both datasets (see Methods). This identified 108 high-confidence splicing events in 105 genes (Fig. 1a and Supplementary Table 1). Remarkably, 24 genes encode proteins that have direct or indirect interaction(s) with another protein in the list, thus identifying 11 protein complexes (Fig. 1b). This high number of complexes is highly significant (p < 2e-5) and shows that B52 tends to co-regulate alternative splicing of genes encoding protein partners. GO term enrichment analysis of the 105 genes identifies “regulation of organ growth” and “positive regulation of growth” as the most enriched GO terms (Fig. 1c). Among them, three genes linked to Hippo pathway were identified: *yorkie* (*yki*), *WW domain binding protein 2* (*Wbp2*) and *Zyxin* (*Zyx*). RNA-seq data indicate that B52 depletion promotes skipping of exon 3 in *yki* mRNA, the inclusion of exons 6 and 7 in *wbp2* and the retention of last intron in *Zyx* (Supplementary Fig. 1). The functional consequences of these alternative splicing events are unknown. We focused our study on *yki* which encodes the effector of Hippo pathway.

**Fig. 1.**
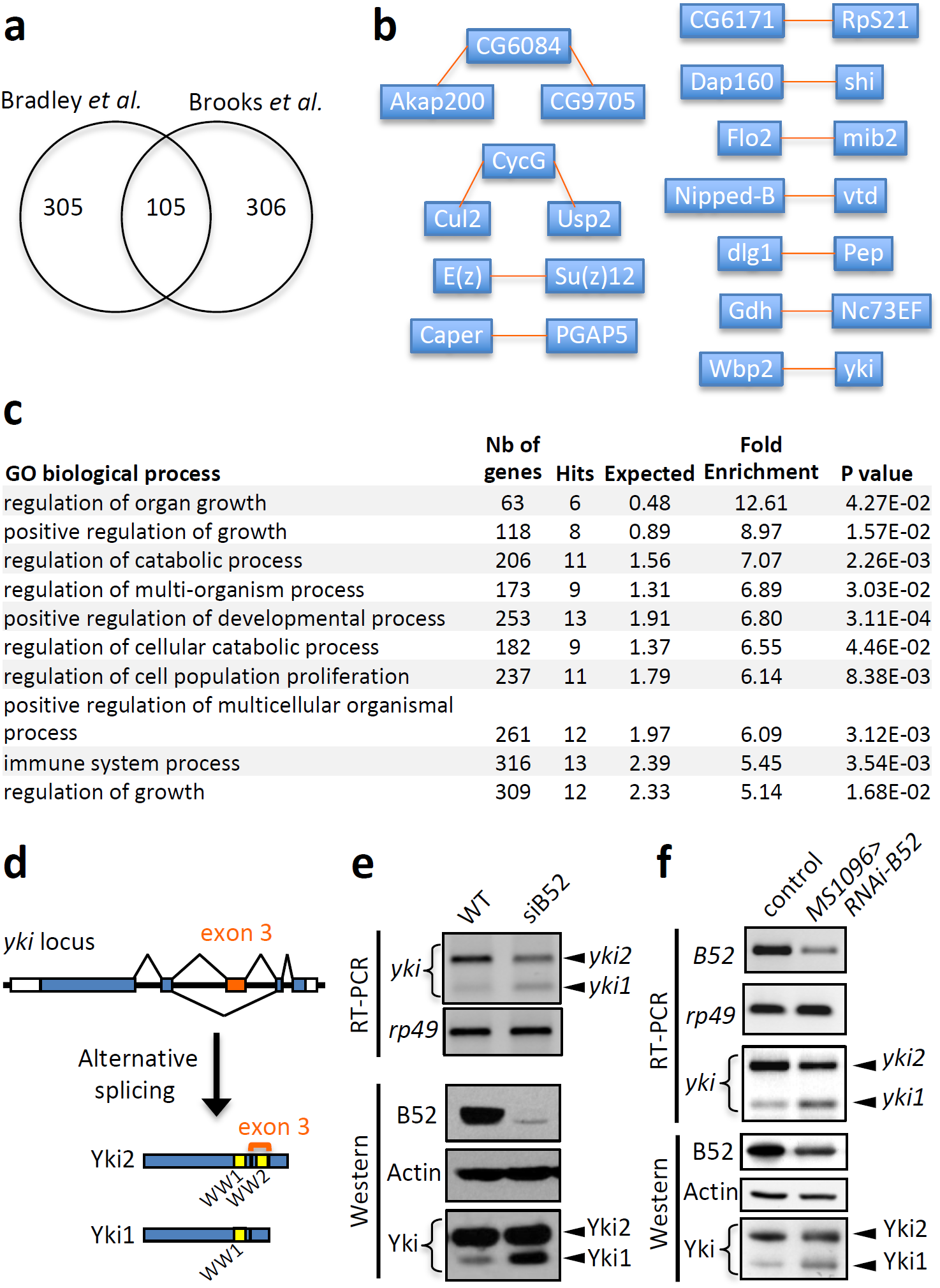
Identification of *yki* as a B52-regulated alternative splicing target. **a**, Number of genes identified by MAJIQ that display 20% or more variation in splicing between wild-type and B52 depleted cells, in each dataset. We identified 108 shared alternative splicing events, corresponding to 105 genes. A comprehensive list of the 108 events is given in Supplementary Table 1. **b**, High confidence protein-protein complexes identified by MIST (Molecular Interaction Search Tool) within the 105 genes identified in **a**. Interactions between proteins can be direct or indirect within the complexes. **c**, GO term enrichment analysis of the 105 genes identified in **a**, performed by PANTHER with Bonferroni correction for multiple testing. **d**, Drawing of *yki* locus and its two isoforms: Yki2 (includes exon 3 and contains 2 WW domains) and Yki1 (skips exon 3 and contains 1 WW domain). **e**,**f**, RT-PCR and western blot showing the effect of RNAi-mediated B52 depletion in S2R+ cells (**e**) and in larval wing discs (**f**). Note that for the wing discs, the Gal4 driver *MS1096* is expressed only in the wing pouch of the disc, thus depletion is partial.

The exon 3 of *yki* gene encodes one of the two WW domains of the protein. Alternative inclusion of this exon leads to production of two isoforms that we named Yki2 and Yki1 according to their number of WW domains (Fig. 1d). We confirmed by RT-PCR and western blotting that RNAi-induced depletion of B52 in S2R+ cells, or in larval wing discs, induces skipping of *yki* exon 3 and increases expression of Yki1 at the expense of Yki2 isoform (Fig. 1e,f). These results identify B52 as necessary for inclusion of *yki* exon 3 and expression of Yki2 isoform.

### B52 depletion reduces growth and Yki activity

To test whether B52 depletion affects Yki activity, we monitored the expression of two reporter genes, *ex-lacZ* and *diap-lacZ*, which are direct targets of Yki. Expression of B52 RNAi in the posterior domain of the wing disc decreases the expression of the two reporter genes in this domain, reflecting a reduction of Yki activity (Fig. 2a). This goes along with a reduction of posterior domain size. These flies are not viable at 25° but are partially viable at 18°C and show a net reduction of wing posterior domain size (Fig. 2b). Using this phenotype, we tested whether *B52* interacts genetically with the Hippo pathway. Overexpression of Yki *(UAS-Yki-V5*, corresponding to long isoform ^15^), as well as depletion of the kinases Hpo or Wts by RNAi, partially rescues the growth defect induced by B52 depletion (Fig. 2b,c). Therefore, increasing Yki activity antagonizes the effect of B52 depletion. Nevertheless, posterior domain size in these contexts remains smaller than the corresponding controls (overexpression of Yki or depletion of Wts or Hpo in the absence of B52 depletion, Fig. 2c). This suggests that B52 depletion affects other genes or pathways necessary for efficient growth. Moreover, the strong overgrowth induced by Yki overexpression, or by Hpo or Wts depletion, may compensate a growth defect in B52 depleted cells, unrelated to the Hippo pathway. This is unlikely because depletion of Tgi, a transcriptional repressor interacting with Sd, which induces a very mild increase of posterior domain size on its own, also rescues the growth defect due to B52 depletion (Fig. 2b,c). Altogether these results indicate that B52 depletion reduces growth at least in part through the Hippo pathway and suggest that it lowers Yki activity by modifying alternative splicing of *yki* mRNAs. We therefore explored the functional differences between the two Yki isoforms.

**Fig. 2.**
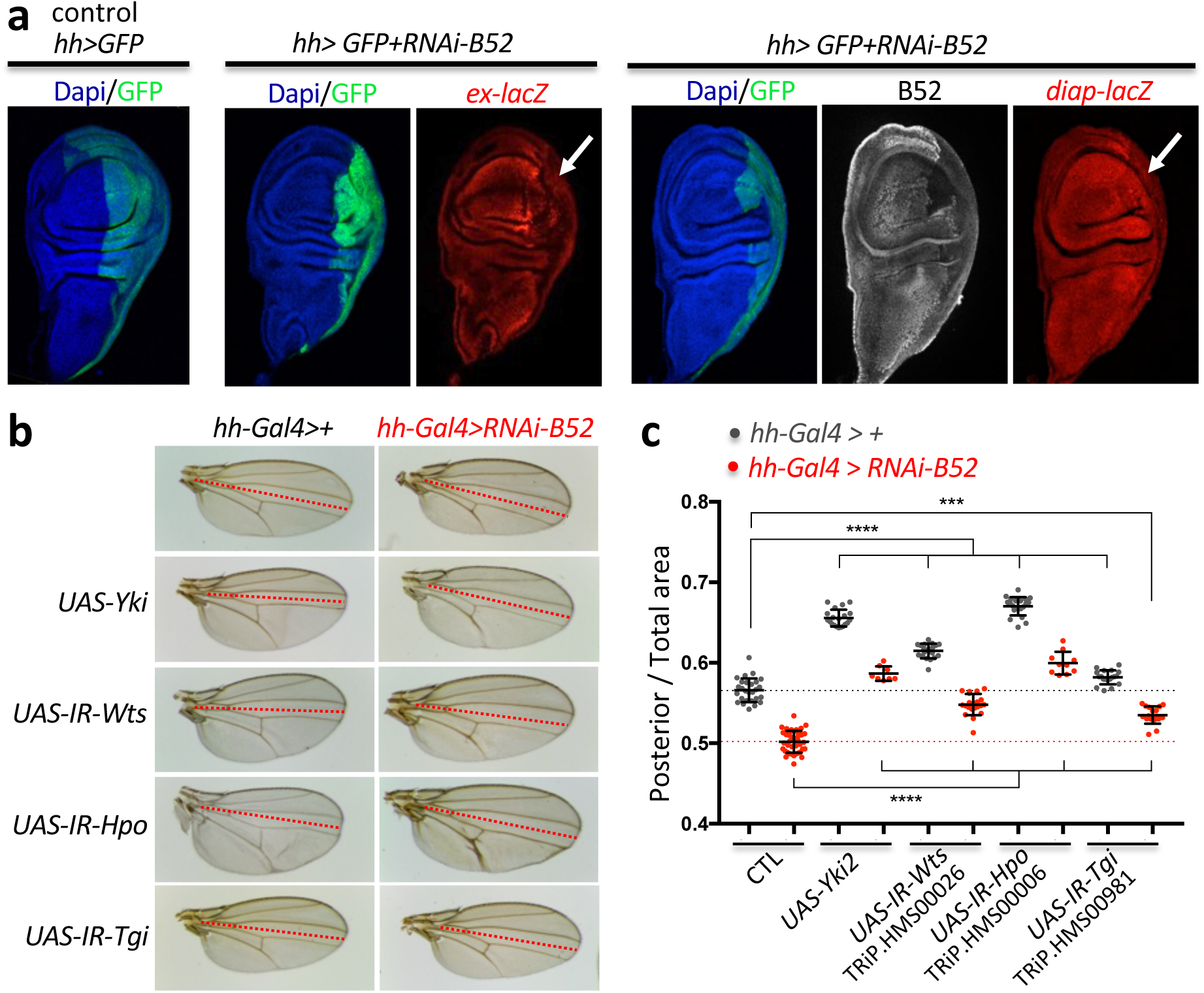
B52 depletion lowers Yki activity and reduces growth. **a**, Expression of *diap-lacZ* and *ex-lacZ* reporters upon depletion of B52 in the posterior part of the wing disc. **b**, Wing phenotype induced by depletion of B52 in the posterior domain of the wing disc (below dotted lines). Flies were grown at 18°C. **c**, Quantification of Posterior/Total wing area in male flies. Each point represents a single wing, bars represent mean with standard deviation. *** p-value<0.001, **** p-value<0.0001 (unpaired two-tailed t-tests). Flies were grown at 18°C.

### Yki1 is a weaker transcriptional activator than Yki2

To compare the activity of Yki isoforms, we created V5-tagged *UAS-Yki1* and *UAS-Yki2* transgenic flies for GAL4-mediated overexpression, by site-specific integration. Compared to Yki2, overexpression of Yki1 isoform induces weaker overgrowth phenotypes in the posterior domain of wings (Fig. 3a) or in the eyes (see Fig. 5a). We analyzed the expression of Yki reporter genes *ex-lacZ* and *diap-lacZ* following Yki isoforms overexpression in the posterior domain of wing discs. Expression of both transgenes is moderately increased by Yki1 overexpression as compared to Yki2 overexpression (Fig. 3b,c). We also analyzed the expression of a *bantam* miRNA sensor, the expression of which is inversely correlated to the level of *ban* miRNA that is a direct target of Yki. Yki1 induces a weaker decrease of *ban*-sensor expression than Yki2 isoform (Fig. 3c). Together these results show that Yki1 has a reduced activity compared to Yki2 *in vivo*.

**Fig. 3.**
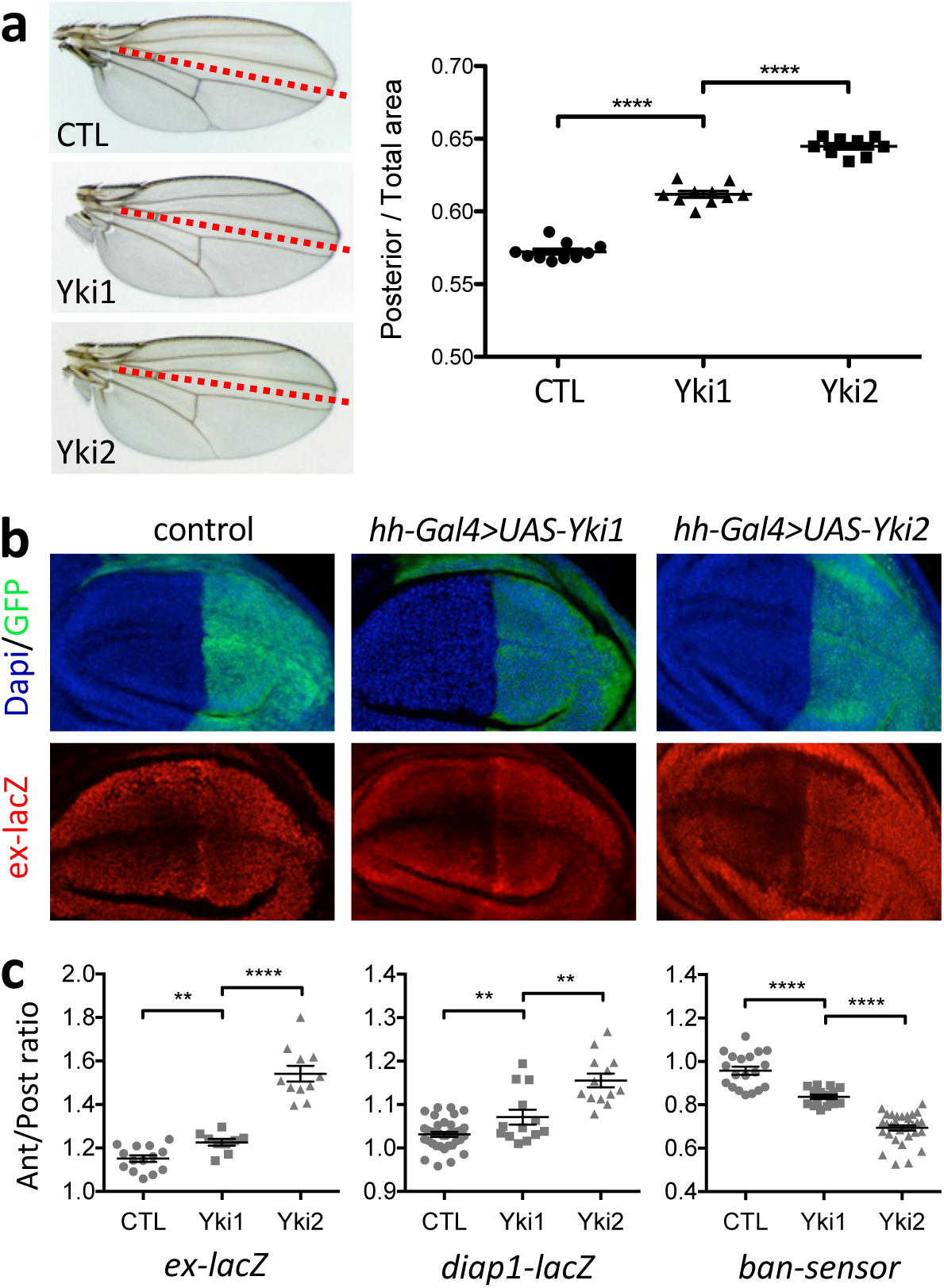
Yki1 isoform is a weaker transcriptional activator than Yki2 isoform. **a**, Phenotype of wings overexpressing Yki1 or Yki2 in the posterior compartment (*hh-Gal4* driver), with the corresponding quantification of the ratio between posterior domain size and total wing size. **b**, Example of immunostaining used to quantify expression of *ex-lacZ* in wild type or upon Yki1 or Yki2 overexpression in wing disc posterior domain. GFP labels the posterior domain. **c**, Quantification of Yki reporter genes expression upon overexpression of Yki isoforms in the wing posterior domain (*hh-Gal4* driver). For each target gene, relative expression between posterior and anterior domain in the wing discs was quantified by immunostaining using anti-ßgal antibody (for *ex-lacZ* and *diap1-lacZ*) or direct visualization of GFP (*ban-sensor*, in this case posterior domain was visualized by immunostaining against V5-tag present in *UAS-Yki* transgenes). In the scatter dot plot, each symbol (circle, square and triangle) represents a single wing disc, bars represent mean with SEM. ** p-value<0.01; **** p-value<0.0001 (unpaired two-tailed t-tests).

**Fig. 4.**
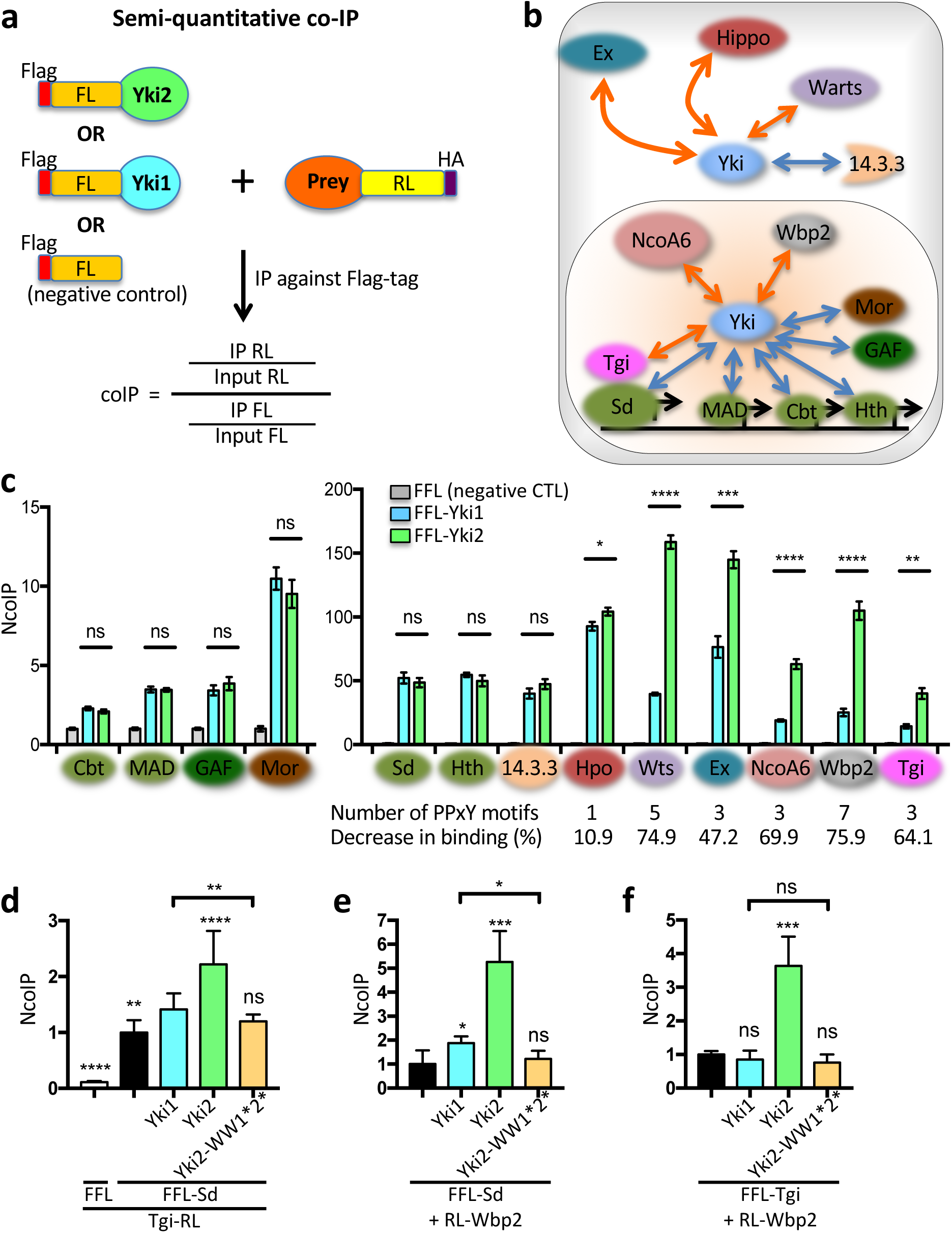
Comparison of Yki isoforms binding properties. **a**, Principle of dual-luciferase co-IP. In all experiments IP is performed against the Flag-tag fused to the Firefly Luciferase (FFL). FFL (unfused to Yki) is used as reference to normalize each interaction. The prey protein is fused to Renilla Luciferase (RL) in C-term for most proteins or in N-term (for Wbp2 and Mor). **b**, Drawing of the monitored protein-protein interactions. Interactions previously shown to involve WW domains are indicated by orange arrows, whereas other interactions are schematized by blue arrows. **c**, Comparison of Yki isoforms binding capabilities by dual-luciferase co-IP. Graphs show the normalized co-IP (NcoIP) results for 13 proteins tested against the two Yki isoforms and the negative control (Flag-Firefly). **d**, Interaction between Sd and Tgi in absence or presence of Yki isoforms. **e**, Interaction between Sd and Wbp2 in absence or presence of Yki isoforms. **f**, Interaction between Tgi and Wbp2 in absence or presence of Yki isoforms.

**Fig. 5.**
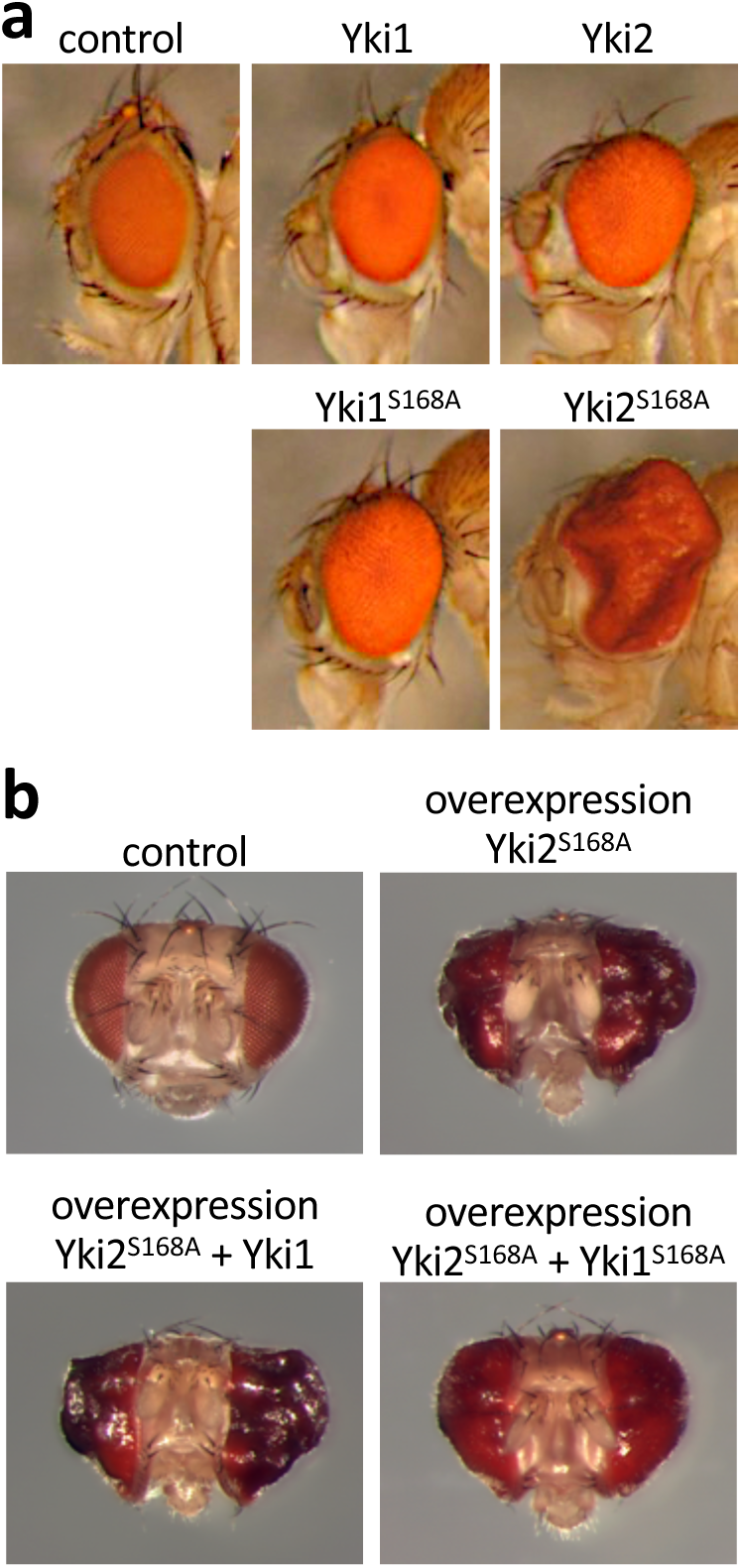
Yki1 isoform can compete with Yki2 isoform in the nucleus. **a**, Phenotypes induced by overexpression of Yki isoforms in the eye under the control of the *GMR-Gal4* driver. **b**, Co-overexpression of Yki isoforms using *GMR-Gal4* driver.

### Absence of the second WW domain modifies the binding and bridging activities of Yki

Yki1 and Yki2 isoforms differ respectively by the absence or presence of the alternative exon 3, which includes the second WW domain. Numerous studies have demonstrated that, in the context of Yki2 isoform, WW domains are required for interaction with several partners containing PPxY motifs such as Ex, Hpo, Wts, NcoA6, Wbp2 and Tgi ^2-6,16-18^. The absence of one WW domain in Yki1 could reduce interaction with such partners. Moreover, binding of proteins that interact with the N-terminal part of Yki, such as Sd or 14.3.3, may also be affected if the structure of the protein is modified by the absence of the domains encoded by exon 3. It is therefore important to address these interactions in the context of full-length proteins and with a quantitative assay. To this end, we developed a dual-luciferase co-Immunoprecipitation (co-IP) method in S2R+ cells inspired by the DULIP method ^19^ described in mammalian cells. Our assay monitors co-IP between a bait protein fused to Flag-tagged-Firefly Luciferase (FFL) and a prey protein fused to Renilla Luciferase (RL). Quantification of luciferases activities ratio after IP with anti-Flag antibodies gives a rapid and sensitive readout of the interaction between the bait and the prey (Fig. 4a). FFL-Yki1 and FFL-Yki2 were used as baits for interactions with thirteen known Yki partners fused to RL as preys (Fig. 4b). We used FFL as a control for non-specific interactions and quantified a normalized co-IP interaction (NcoIP, see methods). By this approach, we detected significant interactions between Yki isoforms and all proteins tested, from modest co-IP with GAF, MAD and Cbt, to high and very high interactions for the other proteins. We found that both isoforms Yki1 and Yki2 interact similarly with the transcription factors Sd, MAD, Cbt and Hth, the chromatin-associated factors GAF and Mor, and with 14.3.3 protein (Fig. 4c). These results show that interactions with these partners are not influenced by the domains encoded by *yki* exon 3. On the other hand, Yki1’s interaction with Hpo, Wts, Ex, NcoA6, Wbp2 and Tgi, which all contain PPxY motif(s), is reduced compared to Yki2 (Fig. 4c). Interestingly, the strength of this decrease correlates with the number of PPxY motifs present in these proteins (Fig. 4c). This is in agreement with *in vitro* studies reporting cooperative binding between multiple PPxY motifs and tandem WW domains of YAP and Yki ^20,21^. Altogether, these results indicate that Yki isoforms interact similarly with transcription factors but that Yki1 cannot efficiently recruit co-activators like NcoA6 or Wbp2 to stimulate transcription.

In addition to increasing the binding of proteins with multiple PPxY motifs, the presence of two WW domains may also allow Yki2 to interact simultaneously with two PPxY-containing proteins. To test this hypothesis, we compared the ability of Yki isoforms to bridge different proteins, by monitoring their co-IP using our assay. Previous reports showed that Yki binds to Sd through its N-terminal part and to Tgi *via* the WW domains ^22,23^. Moreover, Sd and Tgi were shown to interact directly ^4,5^. Yki was shown to compete with Tgi for Sd binding ^4,5^, but was also found in a trimeric complex with Sd and Tgi ^4^. We, therefore, monitored Sd/Tgi interaction in absence or presence of Yki isoforms in transfected cells (Fig. 4d). Using FFL-Sd as bait, we detected co-IP between FFL-Sd and Tgi-RL in the absence of Yki. Both Yki1 and Yki2 isoforms enhanced the co-IP between Sd and Tgi, with Yki2 having a stronger effect. Point mutations in the two WW domains of Yki2 abolish this effect. These results are in agreement with the existence of a trimeric complex between Sd, Tgi and one isoform of Yki.

To further test Yki’s bridging activity, we analyzed the co-IP between FFL-Sd and RL-Wbp2 by the same approach. The co-transfection of Yki1 or Yki2 increases the pull-down between Sd and Wbp2, with Yki2 having a stronger effect. Mutation of both WW domains in Yki2 abrogates this effect (Fig. 4e). These results indicate that, in interaction with Sd, both Yki1 and Yki2 isoforms participate in the recruitment of Tgi or Wbp2, with Yki2 being more efficient owing to its two WW domains. Finally, we tested if the two WW domains of Yki2 can interact with two different proteins carrying PPxY motifs. To this end, we analyzed the co-IP between Tgi and Wbp2 using FFL-Tgi as bait and RL-Wbp2 as prey. Remarkably, Yki2 isoform allowed to pull-down Wbp2 with Tgi, whereas Yki1 isoform did not. As expected, mutation of the two WW domains in Yki2 abrogates the co-IP (Fig. 4f). Altogether these results show that the presence of the second WW domain in Yki2 isoform increases the interactions with PPxY-motifs containing proteins and allows Yki2 to bridge two different proteins with such motifs.

### Yki isoforms can compete with each other in the nucleus

Our results show that, compared to Yki2 isoform, Yki1 isoform interacts similarly with the transcription factors, but less with the transcription coactivators NcoA6 and Wbp2, providing an explanation to its lower transcriptional activity. This suggests that Yki isoforms may compete with each other for binding to the transcription factors. To directly test whether Yki1 isoform can compete with Yki2 isoform *in vivo*, we co-overexpressed them in the eye. We used wild-type forms of Yki isoforms as well as activated versions of these isoforms, carrying the mutation S168A that kills a Wts phosphorylation site and lowers Yki cytoplasmic retention by 14.3.3 proteins ^24^. We verified that this mutation favors the nuclear accumulation of both Yki1^S^168^A^ and Yki2^S^168^A^ isoforms (Supplementary Fig. 2) and increases the overgrowth phenotypes induced by overexpression of these isoforms in the eye (Fig. 5a). Of note, activated Yki1^S^168^A^ induces dramatically weaker overgrowth compared to Yki2^S^168^A^ confirming its reduced transcriptional activity. Overexpression of Yki2^S^168^A^ in the eye induces strong over-growth and remarkably, co-overexpression of activated Yki1^S^168^A^, but not non-activated form Yki1, reduces this phenotype (Fig. 5b), providing evidence of competition between the two isoforms in the nucleus. Altogether, our results are in agreement with a competition model between a fully active isoform Yki2 and a less-active isoform Yki1.

### B52’s effect on growth involves *yki* alternative splicing

Our results show that B52 depletion reduces growth (Fig. 2) and promotes expression of Yki1 isoform through modulation of *yki* alternative splicing (Fig. 1). To directly determine if this contributes to the effect of B52 on growth, we abrogated alternative splicing of *yki* exon 3 by creating flies producing exclusively Yki2 isoform. To this end, we edited endogenous *yki* locus to replace the central part of the gene by a portion of *yki2* cDNA, therefore eliminating the introns surrounding exon 3 (Fig. 6a, see strategy in Supplementary Fig. 3). We generated two *yki^2-only^* alleles, *yki*^*2-only-A*^ and *yki*^*2-only-B*^, corresponding to two independent gene conversion events, which show identical phenotypes. Homozygous *yki*^*2-only*^ flies are viable, despite the fact that *yki*^*2-only*^ embryos display about 50% lethality (not shown). We confirmed by western blotting that *yki*^*2-only*^ flies express only Yki2 isoform and do not produce Yki1 isoform (Fig. 6b). Remarkably, depletion of B52 in wing posterior domain in *yki*^*2-only*^ background induces a weaker phenotype than in a wild type background (Fig. 6c) indicating that B52’s effect on growth is in part due to modulation of *yki* alternative splicing.

**Fig. 6.**
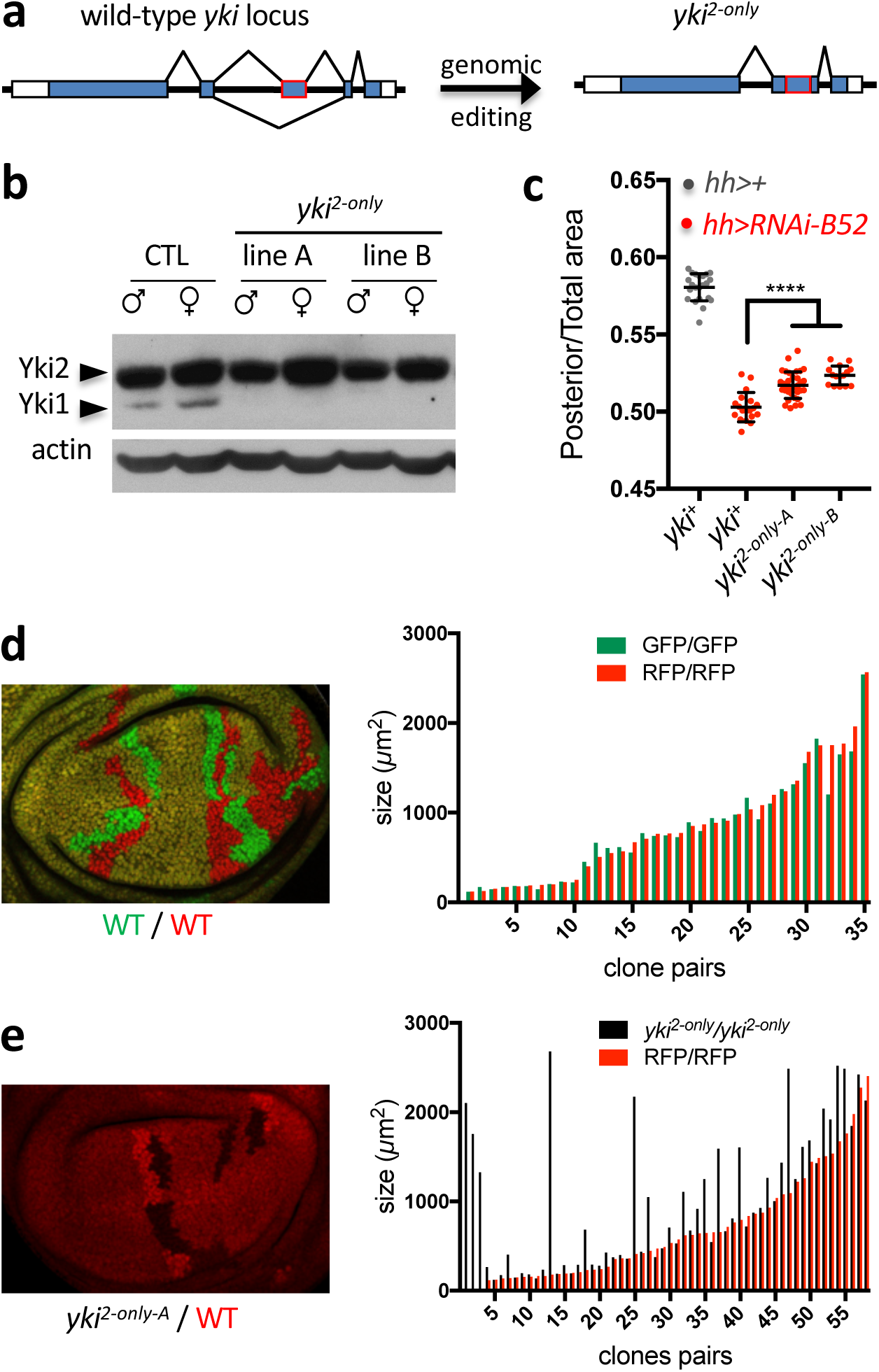
Absence of *yki* alternative splicing reduces the phenotype of B52 depletion in the wing and favors clonal growth. **a**, Structure of the *yki*^*2-only*^ allele compared to wild type *yki* locus. Introns surrounding exon 3 are removed. This allele does not contain any exogenous sequence. **b**, Western blot of adult males and females showing disappearance of Yki1 isoform in the two *yki*^*2-only*^ fly lines. **c**, Comparison of posterior/total area ratio of females wings upon depletion of B52 in a *yki*^*+*^ (wild type) or *yki*^*2-only*^ background. Each point corresponds to a single wing, bars represent mean with SD (t-tests: **** p-value <0.0001). **d**,**e**, Quantification of sibling clones size in the wing pouch. In **d**, sibling wild-type clones are labelled with 2xGFP and 2xRFP. In **e**, unlabelled clones are homozygous for the *yki*^*2-only-A*^ allele whereas sibling clones (RFP/RFP) are wild type. In both graphs, clones pairs are sorted according to RFP/RFP clones size.

Since Yki1 is less active than Yki2 isoform (Fig. 3) and can compete with it (Fig. 5), we asked whether the absence of Yki1 would lead to growth advantage. We did not detect overgrowth in *yki^2- only^* adult flies. Therefore, we addressed if absence of Yki1 isoform influences growth in clonal assays in the wing disc. In this assay, wild-type sibling cells generated by mitotic recombination give rise to clones of similar size (Fig. 6d). On the other hand, homozygous *yki*^*2-only-A*^ cells, obtained by mitotic recombination in heterozygous *yki*^*2-only-A*^*/yki*^*+*^ larvae, often create bigger clones than their sibling wild type *yki*^*+*^*/yki*^*+*^ cells (Fig. 6e). In few cases, *yki*^*+*^*/yki*^*+*^ sibling clones were not recovered, suggesting that these cells were eliminated. These results indicate that the absence of Yki1 in *yki*^*2-only*^ cells increases overall Yki activity and are in agreement with a model of Yki1 isoform acting as a dimmer of Yki activity.

### Alternative splicing of *yki* is required for developmental stability

Despite being viable and fertile, we noticed that several *yki*^*2-only*^ flies in the population display asymmetric wings (Fig. 7a). We measured fly wings areas of wild type and *yki*^*2-only*^ flies, in duplicate to take into account the variability introduced by the manual quantification. This measurement error is low compared to the size difference between right and left wings (Supplementary Fig. 4). The Fig. 7b illustrates the distribution of right and left wing sizes of 40 male and female flies in wild type flies and in the two *yki*^*2-only*^ lines. We noticed an overall reduction of wing size in *yki*^*2-only*^ flies compared to wild type, and an increased difference between right and left sides in several individuals. To quantify the asymmetry between left and right wings, we calculated the fluctuating asymmetry (FA) index FA10 ^25^ which takes into account measurement errors. We observed that *yki*^*2-only*^ flies display higher FA in both males and females than wild type controls (Fig. 7c). The intra-individual variation between the size of right and left wings is recognized as a readout of developmental instability ^26^. These results indicate that *yki* alternative splicing is necessary for proper growth equilibration between sides during development. This is, to our knowledge, the first experimental evidence that alternative splicing participates in the maintenance of developmental stability.

**Fig. 7.**
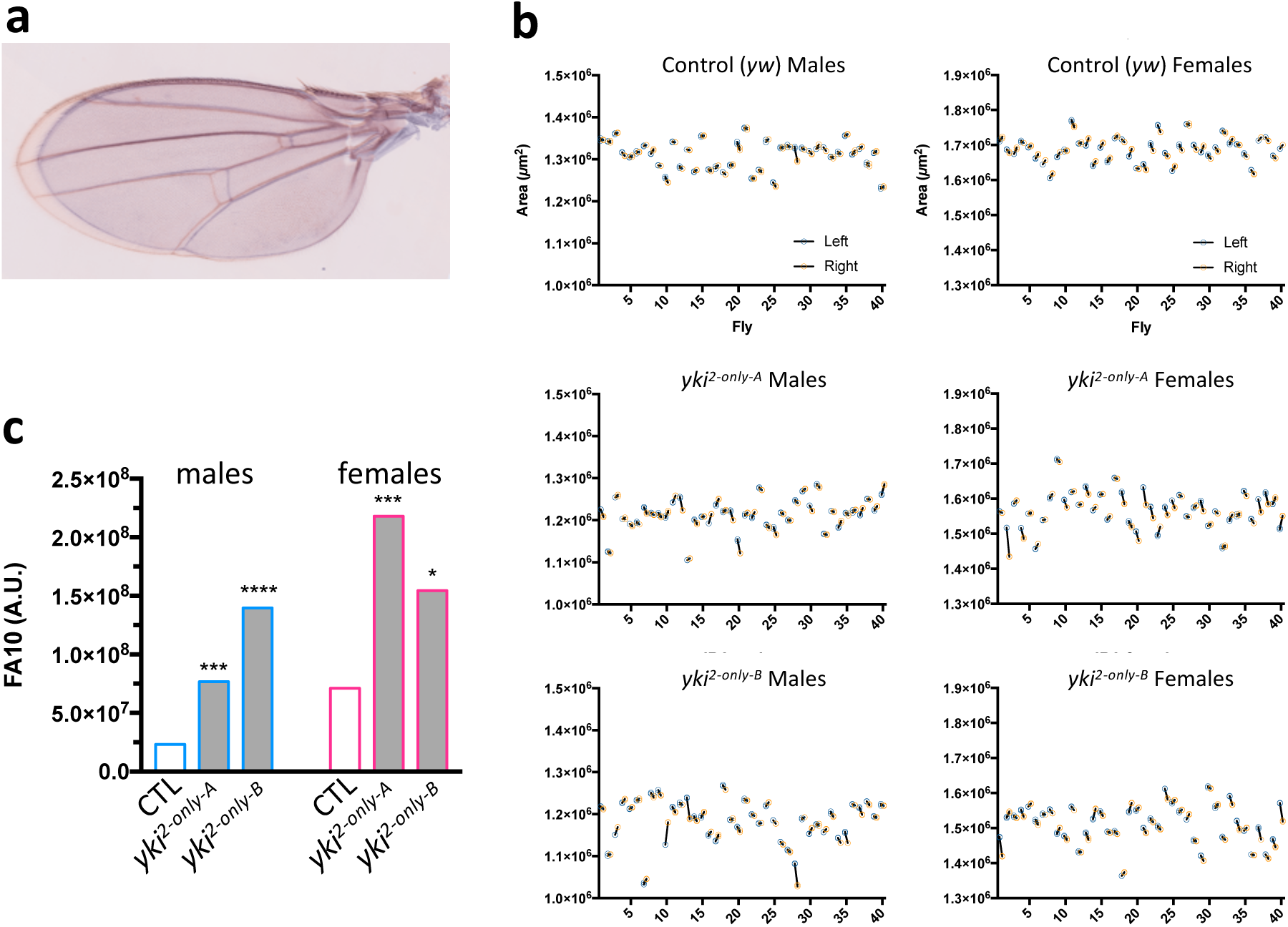
Absence of *yki* alternative splicing increases developmental instability. **a**, Overlay of right and left wings of a *yki*^*2-only-A*^ female showing fluctuating asymmetry. **b**, Distribution of wing size and wing asymmetry among the analyzed flies. Each circle represents the area of a single wing. Each wing was measured in duplicate. The difference between Right (orange) and Left (blue) sides of a single fly is schematized by a black line. **c**, Quantification of fluctuating asymmetry index FA10 in control (*yw*), *yki*^*2-only-A*^ and *yki*^*2-only-B*^ flies (n=40) (F-tests: * p-value<0.05; *** p-value<0.001; **** p-value<0.0001).

## Discussion

Here we show that alternative splicing of *yki* exon 3 represents an additional layer of modulation of Yki activity, that unexpectedly participates in buffering developmental noise. We identify the SR protein B52 as the first modulator of *yki* alternative splicing. Significantly, depletion of B52 reduces growth in the wing, and this phenotype is partially rescued in *yki*^*2-only*^ flies, indicating that B52’s effect on growth is in part mediated by its influence on *yki* alternative splicing. B52 depletion induces skipping of *yki* exon 3 and favors the expression of the Yki1 isoform which is a weaker transcriptional activator, compared to the canonical Yki2 isoform. We demonstrate that Yki isoforms interact similarly with transcription factors but differ by their capacity to bind and bridge PPxY-containing proteins. Indeed, Yki1 isoform, which lacks the second WW domain, has reduced interactions with PPxY-containing proteins and lacks the ability to bridge two proteins containing PPxY-motifs. These observations allow us to propose the following model (Fig. 8). Upon repression of the Hippo pathway, unphosphorylated Yki isoforms enter the nucleus. In a wild type situation (Fig. 8a), Yki2 being more abundant, it joins the transcriptionally repressed Tgi-Sd complex. Within this trimeric complex, Yki2 may engage only one WW domain with Tgi, thus being able to recruit through the second WW domain another partner such as Wbp2 or NcoA6, which finally may, or not, displace Tgi. This could reconcile the observation that Tgi enhances Sd-Yki interaction but decreases distance between Sd and Yki as measured by FRET ^4^ suggesting remodeling of the complex. In a context where B52 is depleted (Fig. 8b), Yki1 isoform level is increased and this isoform competes with Yki2. In a complex with Sd and Tgi, Yki1 would be unable to recruit additional partners and activate transcription, unless Tgi leaves the complex and free the single WW domain. Our results argue that a proper balance between Yki1 and Yki2 isoform is necessary, since abrogation of *yki* alternative splicing increases developmental instability (Fig. 8c). Somehow surprisingly, despite the fact that *yki^2- only^* cells show growth advantage in clonal assays, *yki*^*2-only*^ flies do not show overgrowth of the wings. It has been shown that, in addition to its intrinsic effect on tissue growth, Yki is involved in systemic growth by interacting with ecdysone signaling. Yki is involved in basal expression of ecdysone ^27^, and interacts with the ecdysone receptor coactivator Taiman ^28^. Alteration of ecdysone signaling in *yki^2- only^* flies may contribute to the observed growth defects. In addition, it has recently been shown that Yki controls the expression of the Dilp8 hormone which is involved in inter-organ coordination of growth ^29^. Strong *Dilp8* mutants were shown to display high FA ^30^. Significantly, genomic deletion of *dilp8* Hippo Responsive Element is sufficient to increases FA in flies ^29^ indicating that Yki’s control on *dilp8* is required to minimize developmental variability. It is possible that abrogation of *yki* alternative splicing alters Yki activity and its control on *dilp8* expression, thus triggering instability. Nevertheless, we rarely detect upregulation of *dilp8* expression in *yki*^*2-only*^ clones compared to wild type clones suggesting that this modulation is weak or occurs in only a small fraction of the clones (not shown). It is likely that the effects observed in *yki*^*2-only*^ animals are a combination of local and global perturbations of growth signals. It will be important to determine to which extent *yki* alternative splicing is dynamically regulated during normal growth and regeneration, and which signals control this regulation. The splicing factor B52 is a prime candidate to be involved in this process.

**Fig. 8.**
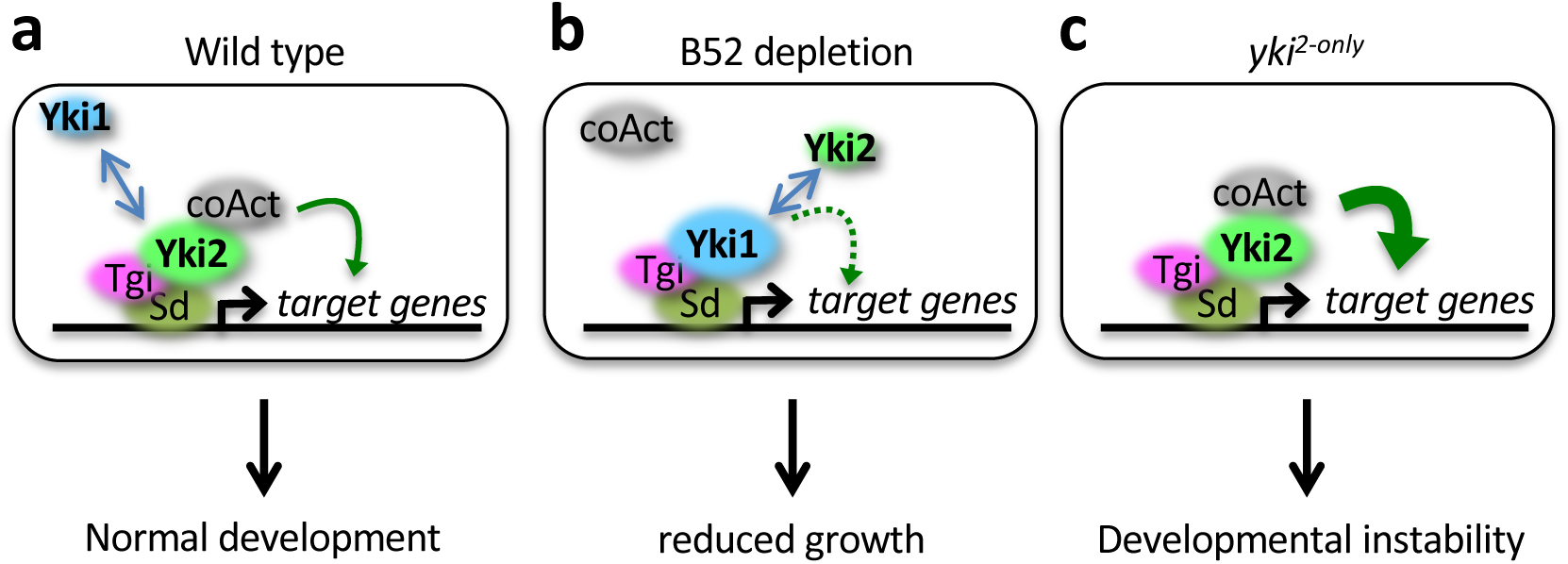
Model of competition between Yki isoforms. See text.

Alternative splicing of pre-messenger RNAs is recognized as a major source of transcriptomic and proteomic diversity ^31,32^. Its extent correlates with organism complexity ^33^ and several examples illustrate that alternative splicing is a source of phenotypic novelty during evolution ^34^. Here we provide an experimental evidence that alternative splicing also participates in the process of canalization, which reflects the resistance of a phenotype to genetic and/or environmental perturbations ^35^. Indeed, abrogating a single alternative splicing event in the genome, in the *yki* gene, is sufficient to increase developmental instability in *Drosophila*. It has been proposed that alternative splicing networks created by some splicing regulators participate in the maintenance of transcriptome stability ^36^. In this view, it is worth noting that B52 tends to control alternative splicing events in genes encoding protein partners. A similar observation was made for the neuronal-specific splicing factor Nova, which modulates alternative splicing of genes encoding protein that interact with one another ^37^. Such co-regulation could be an efficient strategy to control the activity of specific protein complexes and to maintain cell homeostasis. It is therefore tempting to speculate that master regulators of alternative splicing networks also participate to developmental stability.

Our results highlight *yki* alternative splicing as a new level of modulation of Yki activity. It is worth noting that the alternative inclusion of the second WW domain is a conserved feature between *Drosophila* Yki and human YAP, whereas some characteristics of these proteins, such as the presence of a coiled-coiled domain and a PDZ domain in YAP, are not conserved between the two species. The second WW domain of Yki/YAP appears to reside in a separate exon since the emergence of bilaterians ^38^, suggesting that alternative splicing of this exon could be an ancestral mode of modulation of Yki/YAP activity. Analogously to our results in flies, human YAP isoform containing a single WW domain (YAP1) is a weaker transcriptional activator than the YAP2 isoform containing two WW domains ^7^. YAP activity is frequently upregulated in cancer cells ^39^. Therefore, targeting this alternative splicing event to favor skipping of the alternative exon could be a novel strategy to lower YAP activity.

## Methods

### RNA-seq analysis

We used MAJIQ (Modeling Alternative Junction Inclusion Quantification) software (v1.0.7) to identify Local Splicing Variations (LSV) in two previously published datasets from Bradley et al. (2015) (#GSM1552264, #GSM1552267) and Brooks et al. (2015) (#GSM627333, #GSM627334 and #GSM627343). Each dataset contains two replicates of control and B52-depleted cells RNA-seq. The reads were subjected to standard quality control (QC) and filtered according to the following parameters: (1) trimming and cleaning reads that aligned to primers and/or adaptors, (2) reads with over 50% of low-quality bases (quality value ≤15) in one read, and (3) reads with over 10% unknown bases (N bases). We used Trimmomatic software (v0.36) to remove primers and bad quality reads. After filtering, we removed short reads (<36bp), the remaining reads are called “clean reads” and stored as FASTQ format. Reads were aligned to *Drosophila melanogaster* reference genome release 6.19 using STAR (Spliced Transcripts Alignment to a Reference) software (v2.5.4b). STAR outputs were stored in BAM files. BAM files were then submitted to MAJIQ analysis pipeline. LSV definitions were generated and quantified by MAJIQ. MAJIQ applies several normalization factors to the raw values before to compute normalized PSI (Percent Selected Index) and compare them between replicates. To make a selection of best candidate genes we used a ΔPSI threshold of 0.2. All others parameters were used with default values. Gene and PSI lists for each dataset were compared to identify common events between them. This identified 108 alternative splicing events in 105 genes that show reproducible change of alternative splicing upon B52 depletion. GO term analysis was performed with PANTHER (http://pantherdb.org/). Sashimi plots were created with IGV (Integrative Genomics Viewer, https://igv.org/). Protein-protein interactions within the 105 identified genes were tested using Molecular Interaction Search Tool (MIST; http://fgrtools.hms.harvard.edu/MIST/) using the high confidence threshold which retains only the interactions (direct or indirect) supported by several experimental methods ^40^. The number of complexes identified was compared to the distribution of the number of complexes found in 50 sets of 105 random genes, which fits a Poisson distribution (λ=2.36).

### Fly strains and Genetics

*Drosophila* were maintained on standard cornmeal-yeast medium. Experiments were performed at 25°C, except for the analysis of wing phenotype of flies expressing B52 RNAi under the control of *hh- Gal4* that were performed at 18°C, as mentioned in figure legends. Inducible RNAi lines used were *UAS-IR-B52* (GD8690), *UAS-IR-Wts* (GD1563 and TRiP.HMS00026), *UAS-IR-Hpo* (TRiP.HMS00006), *UAS-IR-Tgi* (TRiP.HMS00981) and were previously validated in the literature.

*UAS-Yki1* and *UAS-Yki1*^*S*^*168*^*A*^ transgenes were constructed from *pUAS-Yki-V5-His* and *pUAS-Yki*^*S^168^A*^*-V5-His* clones in *pUAS-attB* vector, kindly provided by K. Irvine (these clones contain Yki2 isoform cDNA). A *Xho*I site located between *UAS* sequences and start codon, flanked by two *EcoR*I sites, was deleted by *EcoR*I digestion and re-ligation of these vectors. This generated *UAS-Yki2-V5-His* and *UAS-Yki2*^*S^168^A*^*-V5-His*. The *Sfi*I–*Xho*I fragment containing the C-terminal part of Yki2 was replaced by the corresponding *Sfi*I–*Xho*I amplified from a *yki1* cDNA obtained by RT-PCR (third instar larvae). This generated *UAS-Yki1-V5-His* and *UAS-Yki1*^*S^168^A*^*-V5-His* transgenes. The four constructs *UAS-Yki2-V5-His, UAS-Yki2*^*S^168^A*^*-V5-His, UAS-Yki1-V5-His* and *UAS-Yki1*^*S^168^A*^*-V5-His* were inserted in *attP2* landing site through PhiC31-mediated integration. Injections were performed by Bestgene Inc. In the text, *UAS-Yki* lines are named without the V5-His tag for simplicity.

For immuno-stainings, the following antibodies were used: mouse anti-ßGal (DSHB), mouse anti-V5 (Invitrogen) and rabbit anti-B52 ^41^, using standard procedures. Images were acquired on a Leica SP5 confocal. To compare expression in anterior *vs* posterior part of wing discs, fluorescence intensity was quantified using OMERO (www.openmicroscopy.org) in an equivalent area in the anterior and posterior domains of the wing pouch, excluding dorso-ventral and anterio-posterior boundaries. For clonal analyses, larvae were heat shocked during 30 min at 37°C, either 48h±4 or 72h±4 after egg laying, and then dissected 48h, 68h or 72h after clone induction. Areas of clones were quantified in the wing pouch with OMERO.

The detail of genotypes used in each Figure is given in Supplementary Table 2.

### *yorkie* gene editing

To edit endogenous *yki* locus by CRISPR/Cas9-mediated homologous recombination, a repair construct corresponding to *yki* gene without introns 2 and 3, and containing a piggyBac insertion in intron 1, was created by cloning of multiple PCR fragments amplified with Q5-Taq polymerase (Biolabs). *yki* gene fragments were amplified from *yw* flies. The piggyBac transposon containing the reporter gene *3xP3-DsRed* (expresses DsRed in the eye), was amplified from *pHD-3xFLAG-ScarlessDsRed* (*http://flycrispr.molbio.wisc.edu/scarless) (DGRC #1367) and cloned into a Dra*I site present in *yki* intron 1, thus creating TTAA sequences at both sides of the transposon that are necessary for its excision. We used two guide RNAs targeting exon 2 and exon 4 of *yki*, cloned into pCFD4 (addgene #49411). The plasmid containing the repair construct was co-injected with the plasmid encoding the two guides into *nos-Cas9* embryos. Injection and screening for positive DsRED flies in the progeny were performed by Bestgene Inc. Two DsRED-positive lines, validated by PCR, were selected for excision of piggyBac transposon using a source of transposase (Bloomington #8285). For each line a single excision event was selected to establish a stock. Two independent *yki*^*2-only*^ alleles were obtained, called *A* and *B*. The entire *yki* locus was sequenced in these lines. We detected polymorphism in introns and silent polymorphism in exons. These differences are present between the *yw* line used to create the *yki*^*2-only*^ construct and the *nos-Cas9* line in which the injections were made. Both *yki*^*2-only*^ lines gave similar results.

### Molecular biology

cDNAs corresponding to MAD, GAF, Sd, Hth, 14.3.3, Hpo and Wbp2 were amplified from third instar larvae RNA by RT-PCR, using a forward primer starting at ATG and reverse primer located just upstream (or sometimes including) the stop codon. Forward primer contains CACC sequence upstream from ATG to orient cloning in pENTR/D-Topo (Invitrogen). All cDNAs were entirely sequenced. Other cDNAs were Cabut (DmCD00765693, DNASU), Tgi (DmCD00765105, DNASU), Mor (Addgene #71048), Ex (kindly provided by N. Tapon). NcoA6 was amplified from a plasmid kindly provided by K. Irvine.

For DUAL-luciferase co-IP, we developed three destination vectors for the Gateway system: pAct-Flag-Firefly-RfA, pAct-HA-Renilla-RfA, pAct-RfB-Renilla-HA. The luciferase-gateway cassettes from mammalian vectors pcDNA-Flag-Firefly-RfA, pcDNA5-HA-Renilla-RfA and pcDNA5-RfB-Renilla-HA (kindly provided by E. Bertrand) were cloned between *Kpn*I and *Nhe*I sites of *pAFW* vector backbone (Drosophila Genomics Resource Center). cDNAs cloned into pENTR/D-Topo were transferred to the appropriate destination vector by LR recombination (Invitrogen). Mutation of the two WW domains were introduced in Yki2 cDNA cloned in pENTR/D-Topo with Q5 Site-Directed Mutagenesis Kit (Biolabs). The sequence WxxP in each WW domain was mutated into FxxA to abolish binding without strongly modifying its structure ^42^.

### Cell culture and co-immunoprecipitations

*Drosophila* S2R+ cells were maintained in Schneider’s medium (Invitrogen) supplemented with 10% fetal bovine serum (Sigma), 50 U/ml penicillin and 50 μg/ml streptomycin (Invitrogen) at 27°C. For B52 depletion, S2R+ cells were treated with a mix of two dsRNA (produced by *in vitro* transcription) targeting exon 2 and exon 9 of *B52*, for 72H. RNAs were extracted with trizol (Sigma). Proteins were prepared in urea buffer (63 mM Tris HCl pH7.5, 2% SDS, 5% 2-mercaptoethanol, 8M urea). Primary antibodies used for western were: rabbit anti-Yki ^43^, rat anti-B52 ^44^, mouse anti-actin (DSHB).

For dual-luciferase co-IP, S2R+ cells were transfected with two plasmids (0.15µg each), in quadruplicate, in 24-wells plates (400000 cells/well) using effectene (Qiagen). Typically, 4 plates were handle at the same time to perform 24 co-IPs in quadruplicate. After 48h of transfection, cells were washed with PBS and lysed with 250µl HNTG buffer (20mM Hepes pH7.9, 150 mM NaCl, 1mM MgCl_2_, 1 mM EDTA, 1% triton, 10% glycerol) supplemented with proteases inhibitors (Halt inhibitor cocktail, Thermo Scientific). Immunoprecipitation (IP) were performed on each lysate in 96-wells Neutravidin plates (Termo Scientific) previously coated for 2h with biotinylated anti-Flag antibody (Bio-M2, Sigma) in HNTG (2 µg antibody/well). IP were performed with 100µl of lysate/well and incubated overnight at 4°. IP were washed 5 times with 200 µl HNTG/well for 5 min at 20° on a thermomixer (Eppendorf) with intermittent shacking. Firefly luciferase (FL) and Renilla luciferase (RL) activities were then quantified with DUAL luciferase reporter assay (Promega) using 50µl of reagents/well and an InfiniteF200 reader (TECAN). To quantify input, 10µl of each lysate were transferred in a white 96 plate and quantified as the same time as IP plate with the same procedure. The level of co-IP is quantified by calculating the level of co-IP normalized to the efficiency of IP: coIP=(RL^IP^/RL^Input^)/(FL^IP^/FL^Input^). For each prey tested, a control co-IP using Flag-Firefly (FFL) as bait was performed and used as reference (interaction set to 1). Each coIP value was normalized to the mean of the quadruplicate control co-IP: NcoIP = coIP^FFL-bait^/MEAN(coIP^FFL^).

For bridging experiments between a FL-tagged bait and a RL-tagged prey, in absence or presence of Yki isoforms, the same procedure was used with a co-transfection of 0.15µg of Firefly plasmid, 0.15µg Renilla plasmid and 0.3µg pAct-Myc-Yki plasmid (or a control plasmid expressing GFP for control without Yki). Transfections and IPs were performed in quadruplicate as described above. Interaction detected in absence of Yki was used as reference to calculate the normalized co-IP ratio NcoIP. In all co-IP, results represent mean ± s.e.m. Student’s t-tests (unpaired two-tailed) were performed using GraphPad Prism and illustrated as: * p-value<0.05, ** p-value<0.01, *** p-value<0.001, **** p-value<0.0001.

### Wing measurements

For measurements of posterior *vs* total area (experiments with *hh-Gal4* driver), young flies (1-3 days) of the appropriate genotypes were stored in isopropanol. Wings were mounted in Euparal (Carl Roth GmbH) on glass slide with coverslip and baked overnight at 65°. Pictures were acquired on a Leica M80 stereomicroscope equipped with a Leica IC80 HD camera using LAS software. Measurements were performed with OMERO (www.openmicroscopy.org).

For quantification of Fluctuating Asymmetry (FA), L1 larvae of the appropriate genotypes were collected 24 to 28h after egg laying and reared at 30 animals per tube at 25°C. Corresponding adult flies were collected, stored in isopropanol and their left and right wings were mounted as pairs. Slides were digitalized using Nanozoomer (Hamamatsu). Measurements of wing areas were performed with ImageJ. Each wing was measured twice in two independent sessions, by one or two persons. In rare case where the variation between replicate measurements was superior to 0.5% of total wing area, the wing was quantified again to minimize measurement error. The FA10 index was used to estimate FA, i.e. FA corrected for measurement error, directional asymmetry and inter-individual variation ^25^. For all genotypes, the interaction individual/side was significant, indicating that FA was larger than measurement error (Supplementary Fig. 4). Conventional two-way mixed model ANOVAs were applied to areas data using GraphPad Prism Software. These values were used to calculate FA10 index: FA10=(MS_Interaction_–MS_Residual_)/2. To compare FA10 values between genotypes, we used F-test to compare variance of the samples.

## Acknowledgements

We thank Nicolas Tapon for helpful comments on the manuscript and Thierry Gostan for his advices on statistical analyses. We thank Kenneth Irvine, Nicolas Tapon, Alexandre Djiane and Edouard Bertrand for sharing flies, constructs, and antibodies. We thank the Montpellier RIO Imaging facility for microscopy. We acknowledge the Bloomington Drosophila Stock Center for providing flies stocks, the Developmental Studies Hybridoma Bank (DSHB) for antibodies, and the Drosophila Genomics Resource Center (DGRC), DNASU and Addgene for plasmids. DS was supported by a PhD fellowship from la Ligue Nationale Contre le Cancer.

## Author contributions

DS and FJ performed genetic experiments and wing quantifications. DS performed immunostainings and associated quantifications. MDT and FJ performed the experiments with S2 cells. LM performed the RNA-seq and splicing analyses. JT financed the project and participated into data analysis. FJ designed the project and wrote the manuscript.

---

**Supplementary Figure 1.**
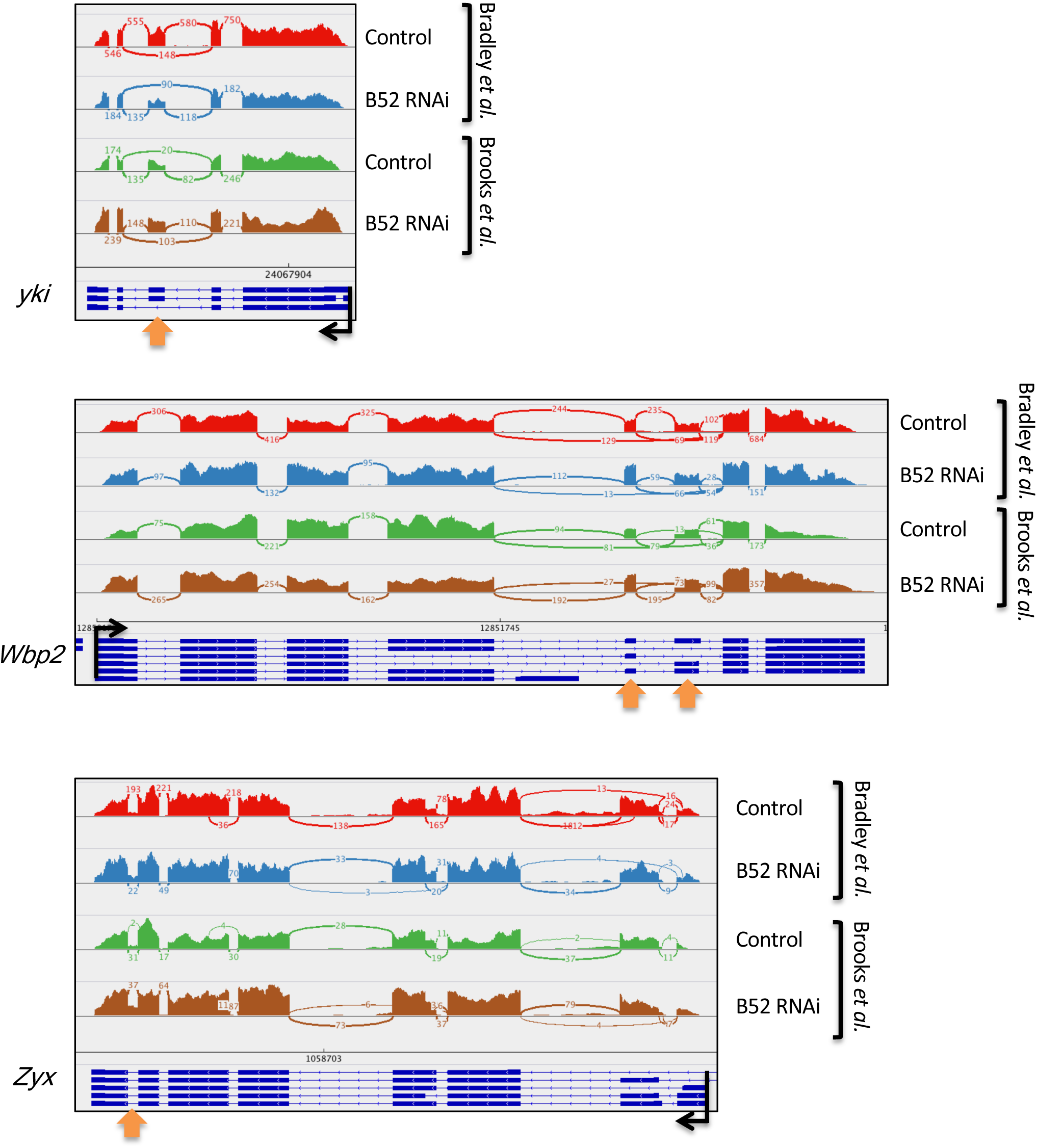
Alternative splicing events modulated by B52 depletion in three genes linked to the Hippo pathway. The Sashimi plots illustrate the alternative splicing variation observed upon B52 depletion in *yorkie* (*yki*), *WW domain-binding protein-2* (*Wbp2*) and *Zyxin* (*Zyx*) in the two datasets. The alternative exons or intron retention are indicated by the orange arrows. Black arrows represent transcription start site.

**Supplementary Figure 2.**
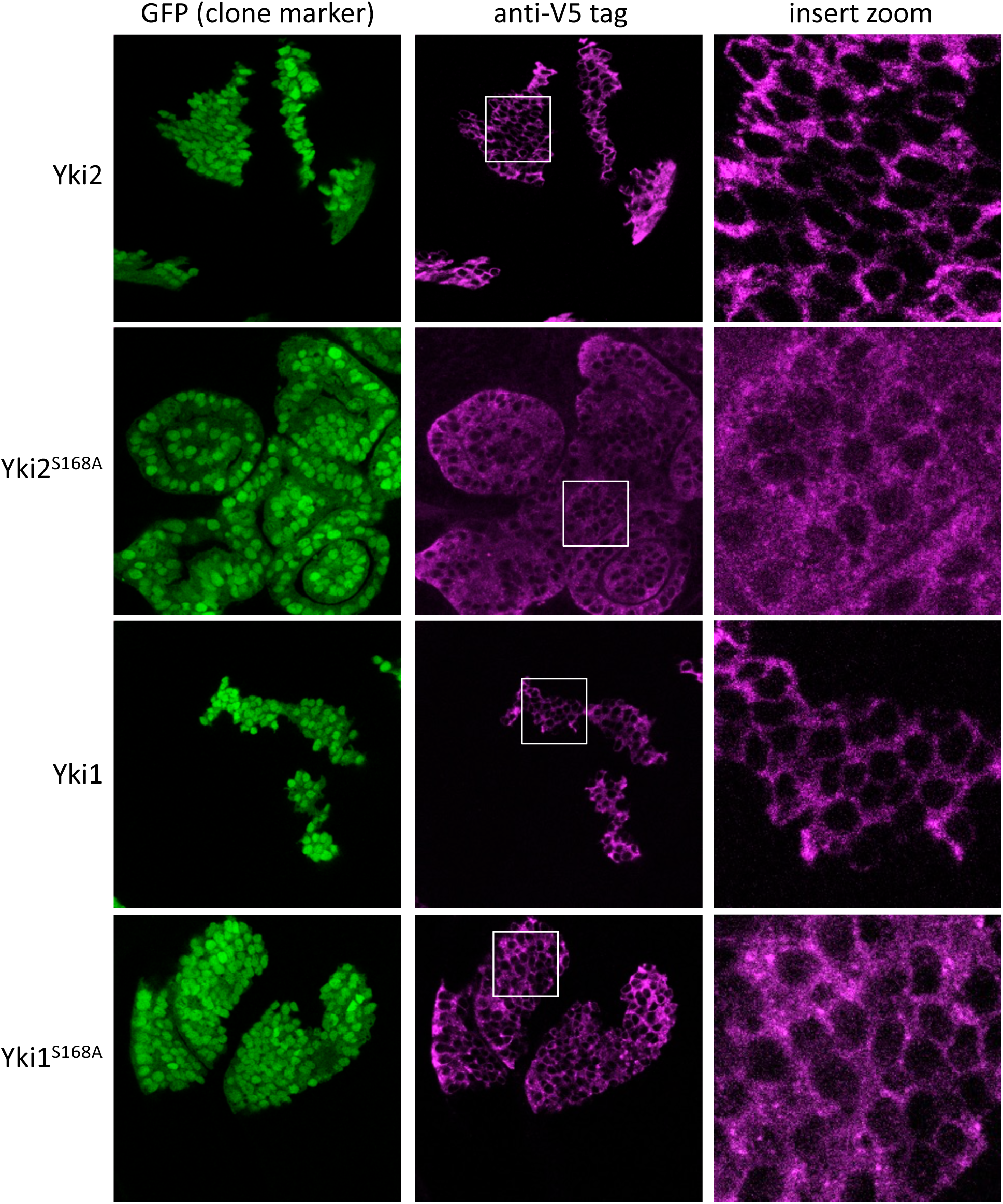
Subcellular localization of Yki isoforms in overexpression clones in wing discs (flip-out clones). Clones are labelled with GFP. Yki proteins are visualized with V5-tag fused to each isoform.

**Supplementary Figure 3.**
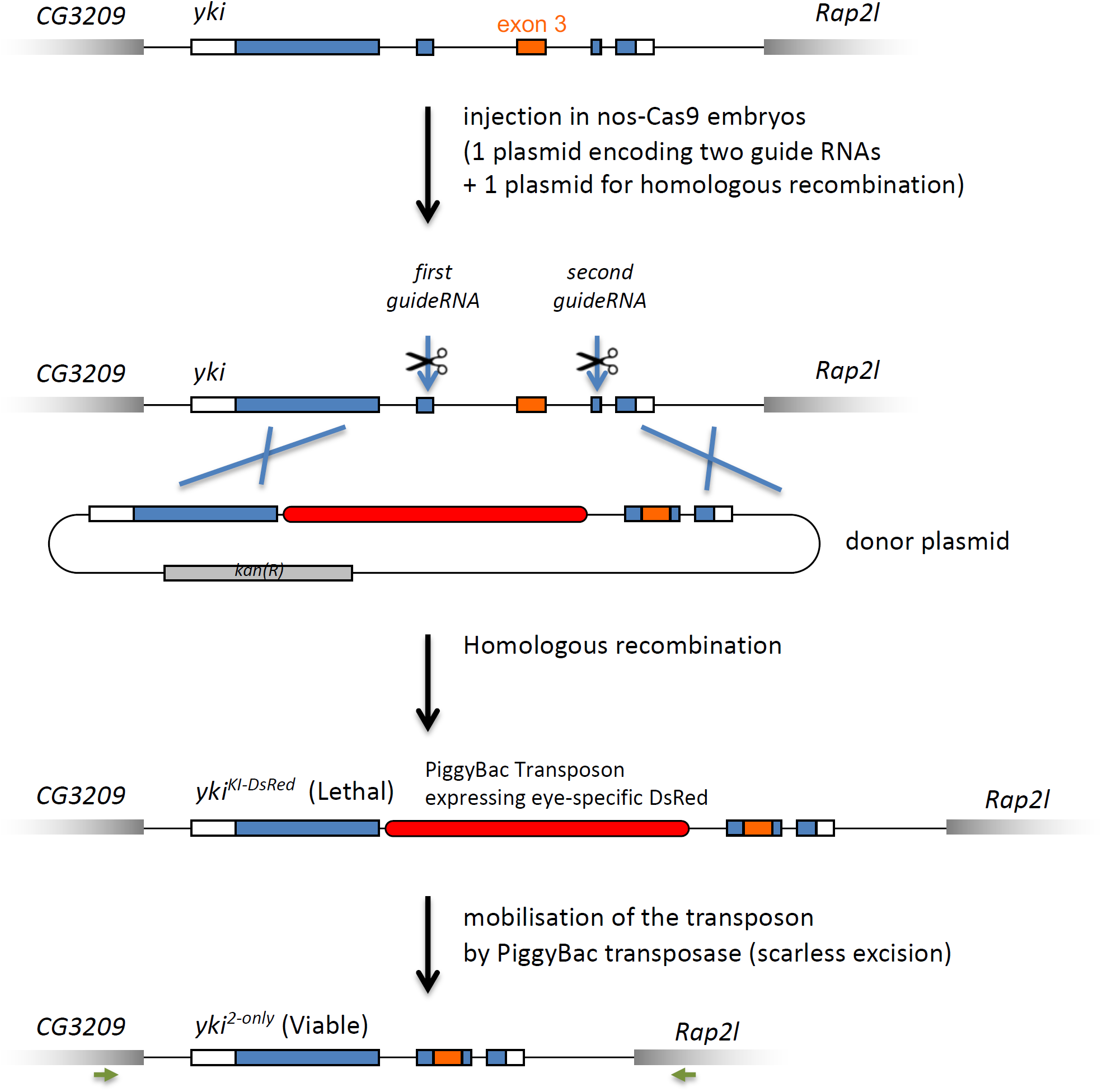
Strategy used to create *yki*^*2-only*^ allele. A donor plasmid was first assembled and cloned in bacteria. It contains the entire *yki* locus deleted for introns 2 and 3 and a PiggyBac transposon, carrying a eye-specific DsRED maker, inserted in the first intron. This construct was used as template for gene conversion after induction of double strand breaks in *yki* locus at the level on exons 2 and 4. Following injection in Cas9-expressing embryos, DsRED positive F1 flies were recovered and analyzed molecularly. Flies carrying the *PiggyBac* insertion in *yki* locus were not viable. Upon excision of the PiggyBac by a transposase provided in *trans*, non-DsRED flies were recovered. These flies were viable and correspond to the *yki*^*2-only*^ allele. Two independent lines (A and B) corresponding to two independent recombination events, were analyzed. The entire locus was sequenced in these lines. We noticed few polymorphism between the two lines, in introns and silent polymorphism in exons. These polymorphisms are present in the parental lines *yw*, used to create the repair construct, and *nos-Cas9*, in which injection was made. The positions of most distal primers used to sequence the locus are shown (green arrows).

**Supplementary Figure 4.**
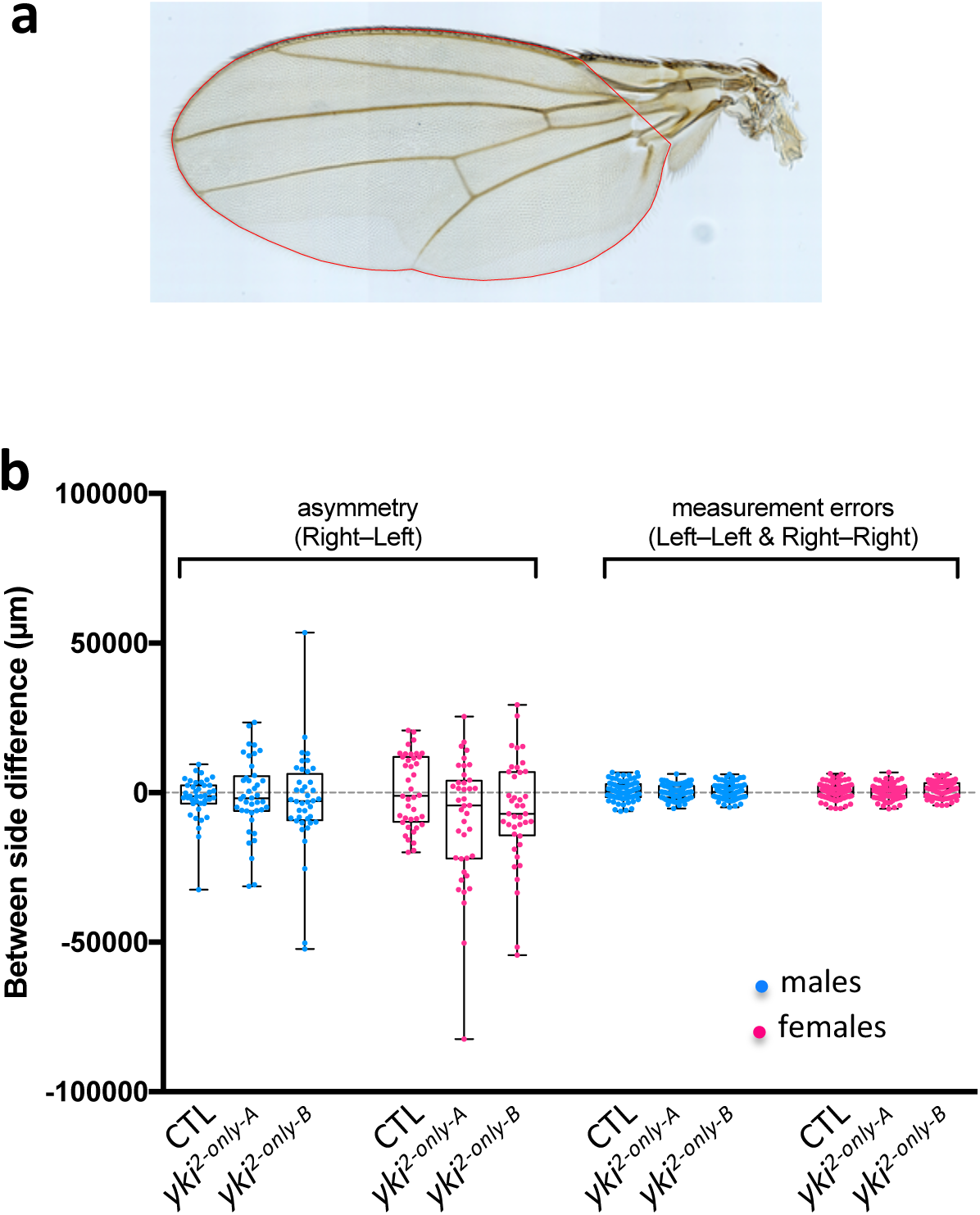
Quantification of fluctuating asymmetry. **a**, Illustration of the domain used to quantify wing area (red line). Each wing was measured twice. **b**, Box plot of Right–Left difference compared to Measurement Errors (corresponding to the difference in replicate measurements). n=40 flies for each genotype.

**Supplementary Table 1.**
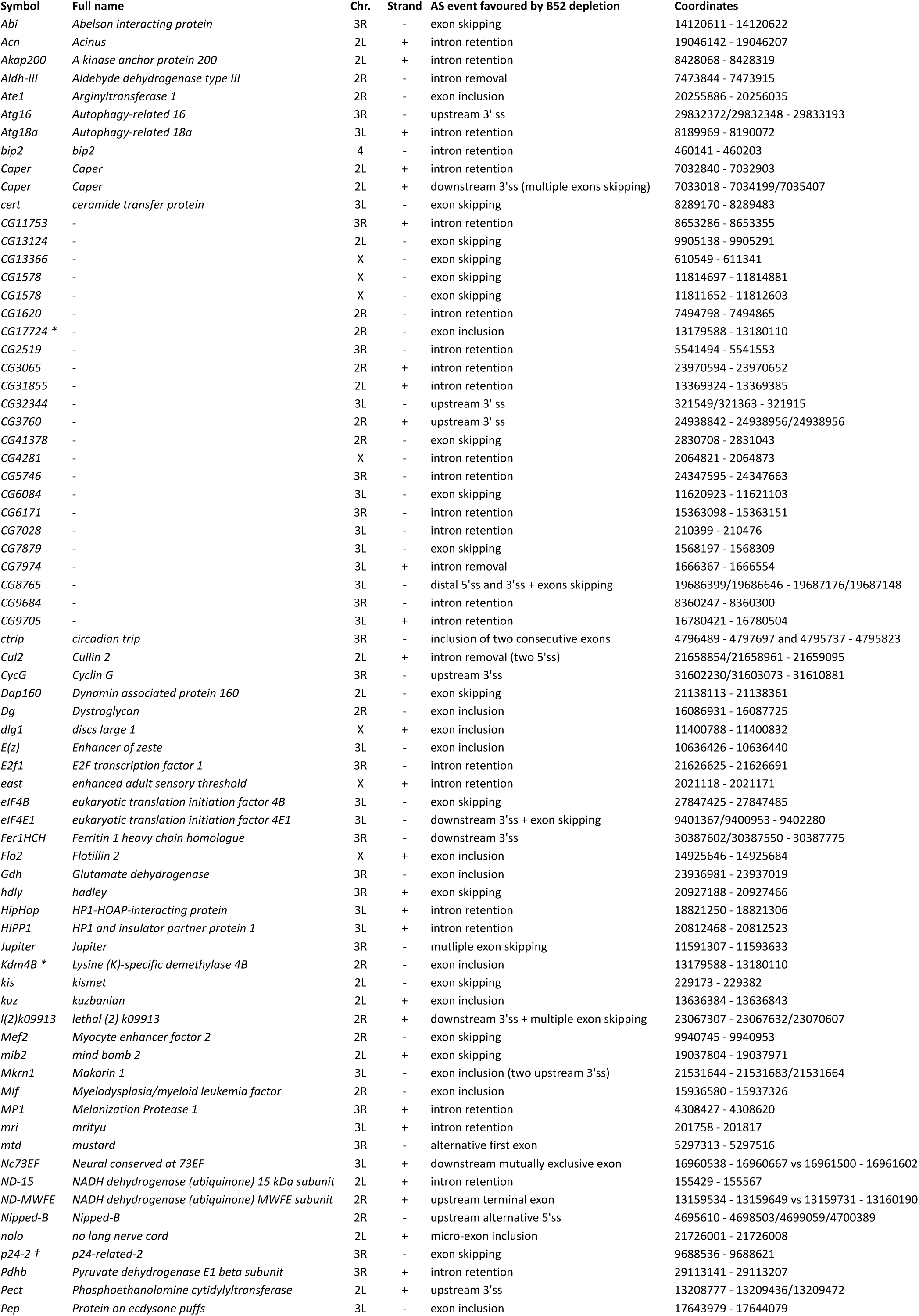
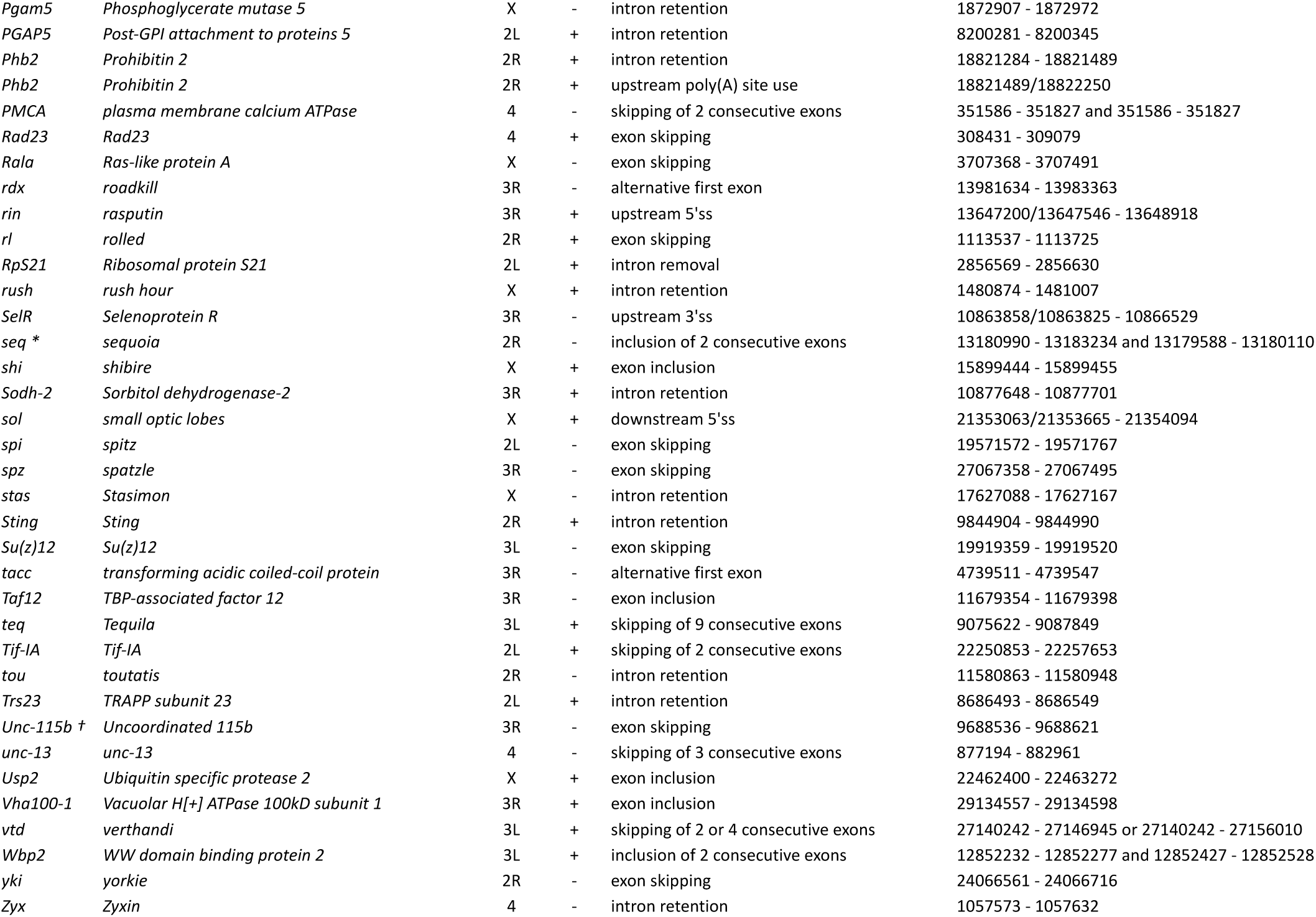
Comprehensive list of Alternative Splicing (AS) events modulated by B52 depletion in S2 cells. AS events were identified based on Local Splicing Variation (LSV) of more than 20% between control cells and B52-depleted cells, in the two datasets of Bradley et al. (2015) and Brooks et al. (2015). Each event was manually annotated to provide a straightforward list of high confidence AS events. The coordinates (based on Dm6.19) correspond to exon boundaries in case of single exon skipping or single exon inclusion, and correspond to intron boundaries in case of intron retention, intron removal, and skipping of multiple exons. Alternative sites are separated by “/”. Symbols * and † denote two groups of overlapping genes containing shared exons.

**Supplementary Table 2.**
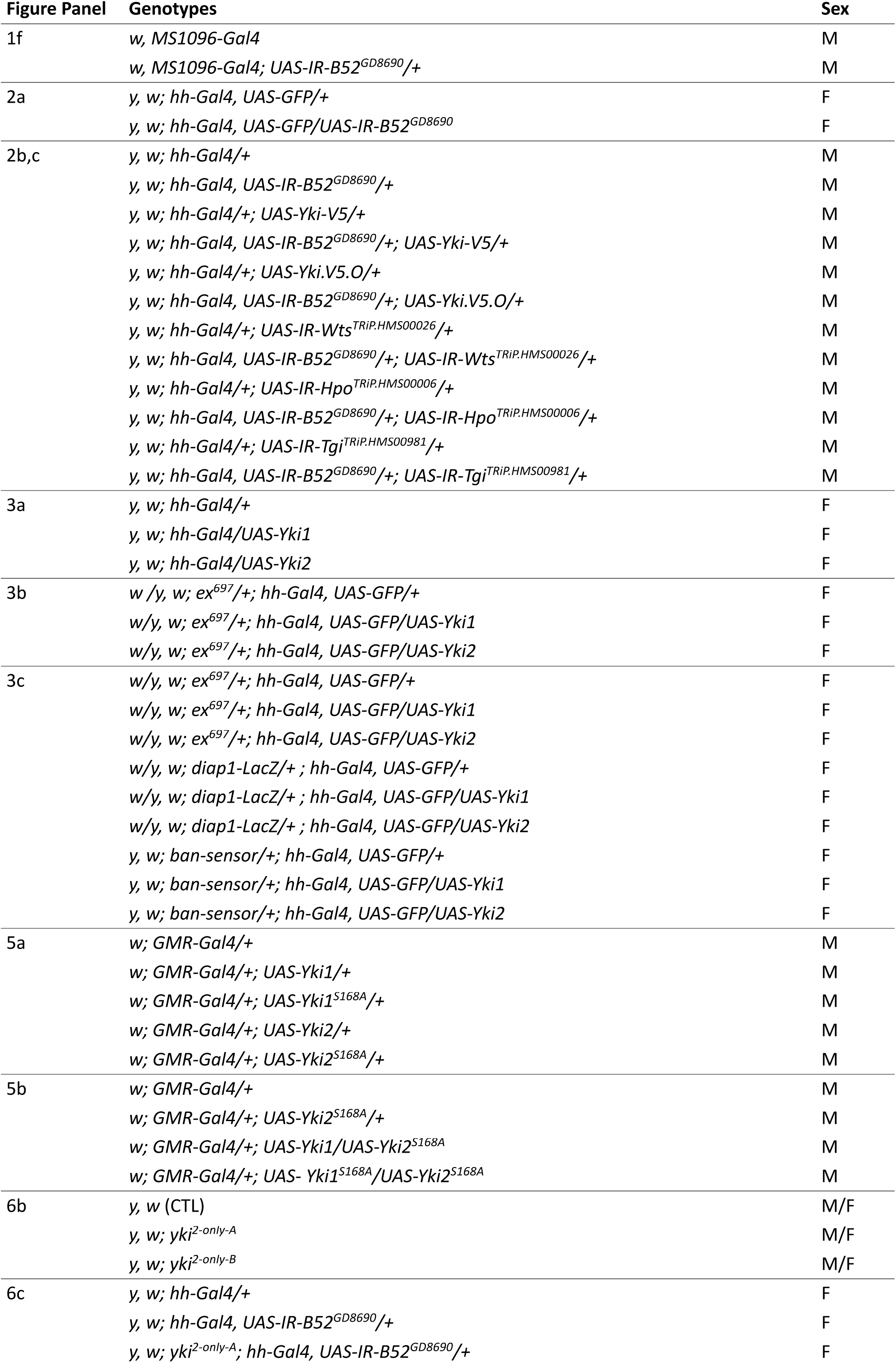
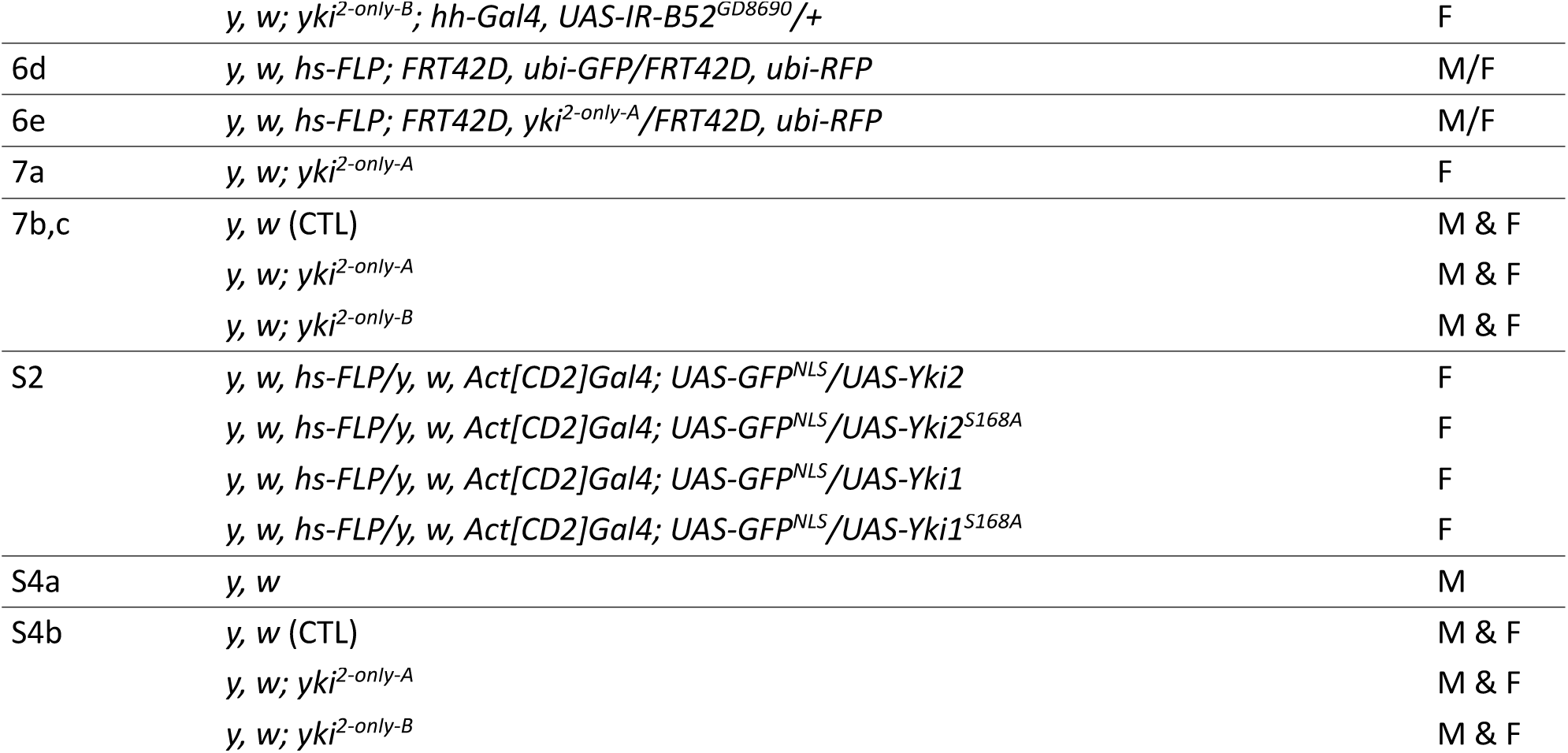
List of genotypes used in each Figure panel.

